# The nervous system is the major target for Gold nanoparticles: Evidence from RNA sequencing data of *C. elegans*

**DOI:** 10.1101/699785

**Authors:** Jihao Mo, Ning Sun, Baolin Yang, Shaoxia Li, Lei Wang

## Abstract

The increasing use of gold nanoparticles (NPs) raises concerns about the potential effect of gold NPs exposure on human health. Therefore, gold NPs exposure is hard to evaluate at the organ level with current measurement technology. The bio-distribution assay showed that intestine was the organ with most gold NPs accumulated in *C. elegans*. However, our data indicated that 62.8% of the significant altered genes were function in the nervous system using tissue enrichment analysis. Notably, developmental stage analysis has demonstrated that NP exposure interfered with the development of animals. Furthermore, the transcription factors DAF-16 was regulating the oxidative stress genes induced by gold NPs. Therefore, we proposed the localization of the oxidative stress genes in the neuron cells and how their expression affect neuron communication. Our results demonstrate that the gold NPs-induced oxidative stress affects the nervous system via physical damage to the neurons and disruption of cell-to-cell communication. Future toxicology research on gold NPs should focus on neurons.

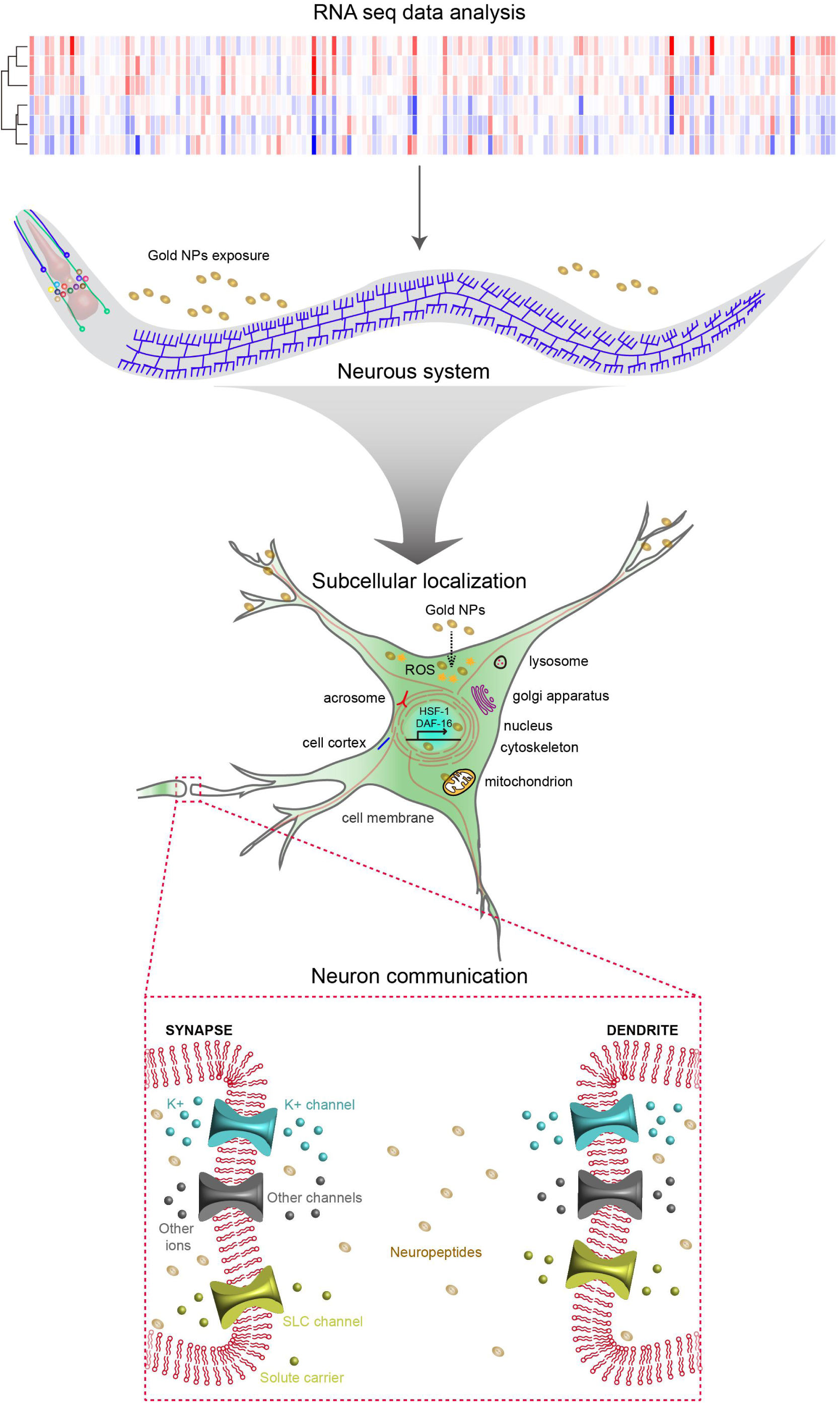
Graphical abstract.

## Introduction

With the development of nanotechnology, there has been growing interest in nano-materials for a wide range of health and medicinal applications. With their nanoscale size, the nano-materials can interact with biological systems at the molecular level to protein, DNA, and RNA ^1,2^. Gold NPs, due to the chemical and physical stability, optical property, like stimuli-response surface-enhanced Raman Scattering, surface plasmon resonance in the near infrared region, is one of the most studied nano-materials in bio-medical applications. In recent decades, the interactions between engineered gold NPs and liver, kidney, blood vessels, tumors have been studied ^3–6^. Additionally, the gold NPs can partially penetrate the blood-brain barrier in the brain, allowing them to act in the central nervous ^7^. Targeted drug delivery using gold NPs has been tested on tumors, the liver, lung, and inhalation ^8–11^. Although relatively low toxicity has been reported on gold NPs, most researchers agree that exposure to nano-scale materials can lead to chronic toxicity, which may be due to the responses at the gene and protein level ^12,13^.

Despite the numerous cytotoxicity studies for gold NPs, which have been investigated *in vivo* and *in vitro*, the results vary based on the type of cells used, the time of exposure, the physical and chemical property and the surface modification of the nanomaterials. *In vitro* studies using human oral cancer cell (HSC-3) have demonstrated that gold NPs enter the cytoplasm, mitochondria, and nucleus of the cells. Few publications claim toxicity of gold NPs at a lower concentration than 1 ppm ^14^. However, when increased to 10 ppm or higher, increased expression of apoptosis, oxidative stress, and pro-inflammatory genes was reported ^15^. Apoptosis, autophagy, DNA damage, and necrosis were also observed in studies in HSC-3, HeLa cells, and liver of BALB/c mice ^16–18^. Furthermore, it has been shown that the higher levels of oxidative stress generated by the large surface of gold NPs, which cause DNA and Mitochondrial damage, further disrupts cellular homeostasis ^17^.

Biodistribution is a key factor to understand how NPs accumulate and interact with each organ. The current quantification techniques require the isolation of each internal organ or the fluorescent/radio imaging modalities to modulate the whole animal. As well as the biodistribution of the NPs, to understand how each organ responds to NPs *in vivo* is a huge challenge in humans but it is possible in small animals. Studies in mice show that 13 nm PEG-coated gold NPs were mainly deposited in the liver, kuffer cells, and the spleen ^19,20^. While other studies in rats proved 18 nm gold NPs were accumulated in the liver and spleen, but 1.4 nm mainly in the kidney ^21^. The intestine of *C. elegans*, have the combined function of liver, spleen and kidney, accumulates most of the gold NPs ^22^. Although these organs accumulate gold NPs, further investigations should be focused on how each organ responds to the gold NPs, and how the expression of genes in each tissue restore homeostasis.

*C. elegans* is a transparent animal with multiple important signaling pathways conserved in mammals, like autophagy, insulin, and apoptosis signaling pathways ^23^. We analyzed the RNA sequencing data set (GSE32521) from gold NPs exposed worm. However, different with the biodistribution data, the results showed that the nerve system was the most affected when exposing organisms to gold NPs. Further studies show that the oxidative stress transcription factor DAF-16 and HSF-1 are the main transcription factors that regulate the gold NP-induced gene expressions. The ion-channel protein analysis indicated that the potassium and solute carrier (SLC) channels are the most altered channels by gold NPs, which could be the direct way of how gold NPs affect neuron activity.

## Materials and Methods

### C. elegans strains and growth conditions

The wild type strain (*N2*) and daf-16 reporter strain (*daf-16* (*zls356*)) were gifted from the Caenorhabditis Genetics Center (St. Paul, MN). The synchronized worms were cultured with nematode growth medium plates. The *E.coli* strain *OP50* was used as a food source. All the experiments were performed at 20°C.

### Measurement of Size distribution and zeta potential

The 40 nm gold NPs were purchased from Biotech (Jieyi, Shanghai). The gold NPs were originally distributed in water, then diluted in S medium. All the samples were sonicated for 30 minutes before the measurement. The size distribution and zeta potential were tested by a Malvern zeta sizer (HTS3000, Germany). The data was collected and graphed by graphpad 7. The data in the manuscript were labeled as average ± standard deviation.

### Measurement of body length and developmental stage

The synchronized wild type nematodes were cultured in NGM plates with 0, 1, 10, 100 mg/L of gold NPs respectively. The *OP50* was added as food resource. The nematodes were observed each day and food was added every 48 hours. Animals were anesthetized by 50 μM sodium azide on slides. The images were taken by Leica microscope with 10X objective lens. The body length was measured by Image J. The developmental stage was determined by observe the phenotype of each organ and the body length. At least 100 worms were counted for each condition. The graph was generated by R. The data in the manuscript were labeled as average ± standard deviation.

### DAF-16 subcellular localization analysis

The DAF-16 subcellular localization was used as a reporter to prove the activation of insulin like receptor pathway. The *daf-16p::daf-16a/b::GFP* allele nematodes were cultured to L4 stage and then transferred to the 60 mm petri dishes with 0, 100 mg/L gold NPs respectively. After 24 hours, the worms were mounted on slides with sodium azide for imaging. The worms of positive control group were exposed to 37°C for 1 hour. The images were taken by Leica fluorescence microscope with a 20X objective lens. The number of cells with clear nucleus localization was counted and plotted by graphpad 7. The data in the graph was labeled as average ± standard error.

### Quantitative Realtime-PCR

The wildtype nematodes were cultured to L4 stage and then transferred to the 60 mm petri dishes with 0, 100 mg/L gold NPs respectively. The RNA of each sample was extracted by TRIZOL (Thermo Fisher, Waltham, MA) as described ^24^. The contaminant DNA was removed by TURBO DNase (Thermo Fisher). The complementary DNA was synthesized from total RNA using Reverse transciptase (Thermo Fisher). QPCR was performed with SYBR Green using a Bio-Rad CFX-96 (Hercules, CA, USA) real-time PCR system. Thirty ng of cDNA was used as templet and 50 nM of paired primer mix of each gene were used for each reaction. Relative mRNA expression of the target genes was normalized to *act-1*. The primers used in this manuscript were listed below. The individual experiment was repeated at least three times. The data in the graph was labeled as average ± standard error.

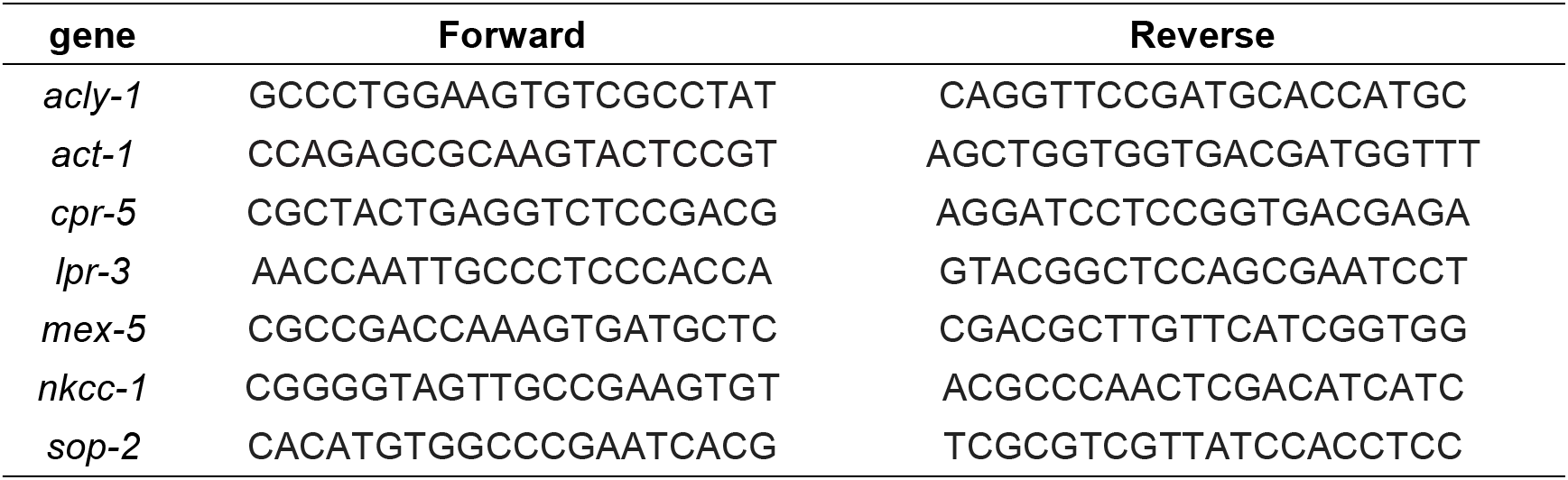

### RNA sequencing data Analysis

The RNA sequencing data, reference series GSE32521, was obtained from Gene Expression Omnibus, National Center for Biotechnology Information. Age-synchronized L3 worms were exposed to 4 nm gold NPs with a concentration of 5.9 mg/L in 50% K-Medium. Then they were cultured with 50% K-Medium and left for another 12h. The RNA was extracted from each of the replicates. Significantly changed genes with 1.5 fold and p < 0.05 and FDR < 0.1 were analyzed ^22^.

### Gene Expression Analysis

The volcano plot was generated by R (version 3.5.2) using package ggplot2. Significantly changed genes with 1.5 fold and *p* < 0.05 were marked red. Genes with 1.5 fold and *p* > 0.05 were marked blue. The tissue enrichment analysis was proceeded by Gene Set Enrichment Analysis function of Wormbase (https://www.wormbase.org/tools/enrichment/tea/tea.cgi) ^25^. The color of each sub-tissue was marked based on the tissue. The motif sequence was analyzed using the motif discovery function of MEME (http://meme-suite.org/tools/meme) ^26^. The length of the enriched motifs was set between 6 to 50 nucleotides.

### Protein function and sub-cellular localization analysis

The protein-protein function analysis was measured using genes that significantly increased in expression. The co-expression, genetic interactions and physical interactions were used as parameters to analyze the interaction using Genemania (https://genemania.org/) ^27^. The top 20 genes with increased expression were marked by differences in function using STRING (https://string-db.org/cgi/input.pl?sessionId=pSsnm1zTtNsB&input_page_show_search=on) ^28^. The sub-cellular localization analysis was investigated based on the structure domain and signal sequence using iLoc-Animal (http://www.jci-bioinfo.cn/iLoc-Animal) ^29^. The number of enriched proteins in each organelle was shown by using a heat map.

### Ion Channel protein Analysis

The proteins specific to each ion channel were isolated from RNA sequencing data as previously reported ^30^. The genes significantly changed in expression, were analyzed and data was shown using a heat map generated by R. The activation of each ion channel was predicted based on the change in expression of ion channel proteins.

## Results

### The nervous system is the most affected by gold NPs

In mice, the liver and spleen are the target organs for gold NPs accumulation ^21^. In *C. elegans*, the intestine plays a role as the liver, kidney, spleen, and stomach, which is also the organ that has most gold NPs accumulated ^22^. For the worms treated with 4 nm gold NPs, 773 genes were significantly increased for at least 2-fold and 268 genes were significantly decreased for at least 2-fold (Fig. 1A). The tissue enrichment assay showed that 62.8% of the significantly altered genes are localized in neuron cells, 14.9% enriched intestine, 7.8% for reproductive system, 1.6% for muscles and 13% for other tissues (Fig. 1B). The PVD neuron and outer labial sensillum have genes significantly increased and decreased, while the thermosensory neuron, amphid sensillum, pharyngeal interneuron, ASE, lateral ganglion, touch receptor neuron, BDU and retrovesicular ganglion have genes only significantly increased, nerve ring and lateral, dorsal nerve cord have genes only significantly decreased (Fig. 1C). The intestine enriched genes show a significant increase only. Meanwhile, the reproductive system and muscle enriched genes shows a significant decrease. From the analysis of the promoter sequence of all the significantly increased genes, a conserved 43 base pairs motif with an e value of 1.5^−222^ was found. This result indicates that the nervous system is the most affected organ by gold NPs exposure and most of the significantly increased genes are enriched in the nervous system and intestine. Most of the significantly decreased genes are enriched in the reproductive system and muscle. To further understand how gold NPs interact with the nervous system, the significantly increased genes which enriched in the nervous system were analyzed by protein-protein interaction assays. In all the significantly increased genes, 20 of them showed more interaction with other proteins (Fig. 1E). Function analysis showed that they are involved in cellular component organization (*let-2*, *dyn-1*, *unc-51*, *unc-54*, *chc-1*, *ama-1*, *sec-24.2*, *eif-3.B*, *tfg-1*, and *F13B9.1*), reproduction *(let-1*, *dyn-1*, *snx-14*, *unc-54*, *H28O16.1*, *dlst-1*, *chc-1*, *copb-1*, *csnk-1*, *eif-3.B*, *ama-1*, *tsr-1*, *sec-24.2*, and *tfg-1*), multicellular organismal development (*let-1*, *dyn-1*, *T22F3.3*, *unc-54*, *H28O16.1*, *dlst-1*, *chc-1*, *csnk-1*, *eif-3.B*, *ama-1*, *tsr-1*, *sec-24.2*, *pme-5*, and *tfg-1*), nematode larval development (*let-2*, *dyn-1*, *unc-51*, *T22F3.3*, *unc-54*, *H28O16.1*, *dlst-1*, *chc-1*, *csnk-1*, *eif-3.B*, *ama-1*, *sec-24.2*, and *tfg-1*) and embryo development in birth or egg hatching (*let-1*, *dyn-1*, *T22F3.3*, *H28O16.1*, *dlst-1*, *chc-1*, *csnk-1*, *eif-3.B*, *ama-1*, *tsr-1*, *sec-24.2*, pme-5 and *tfg-1*) (Fig. 1F).

**Figure 1.**
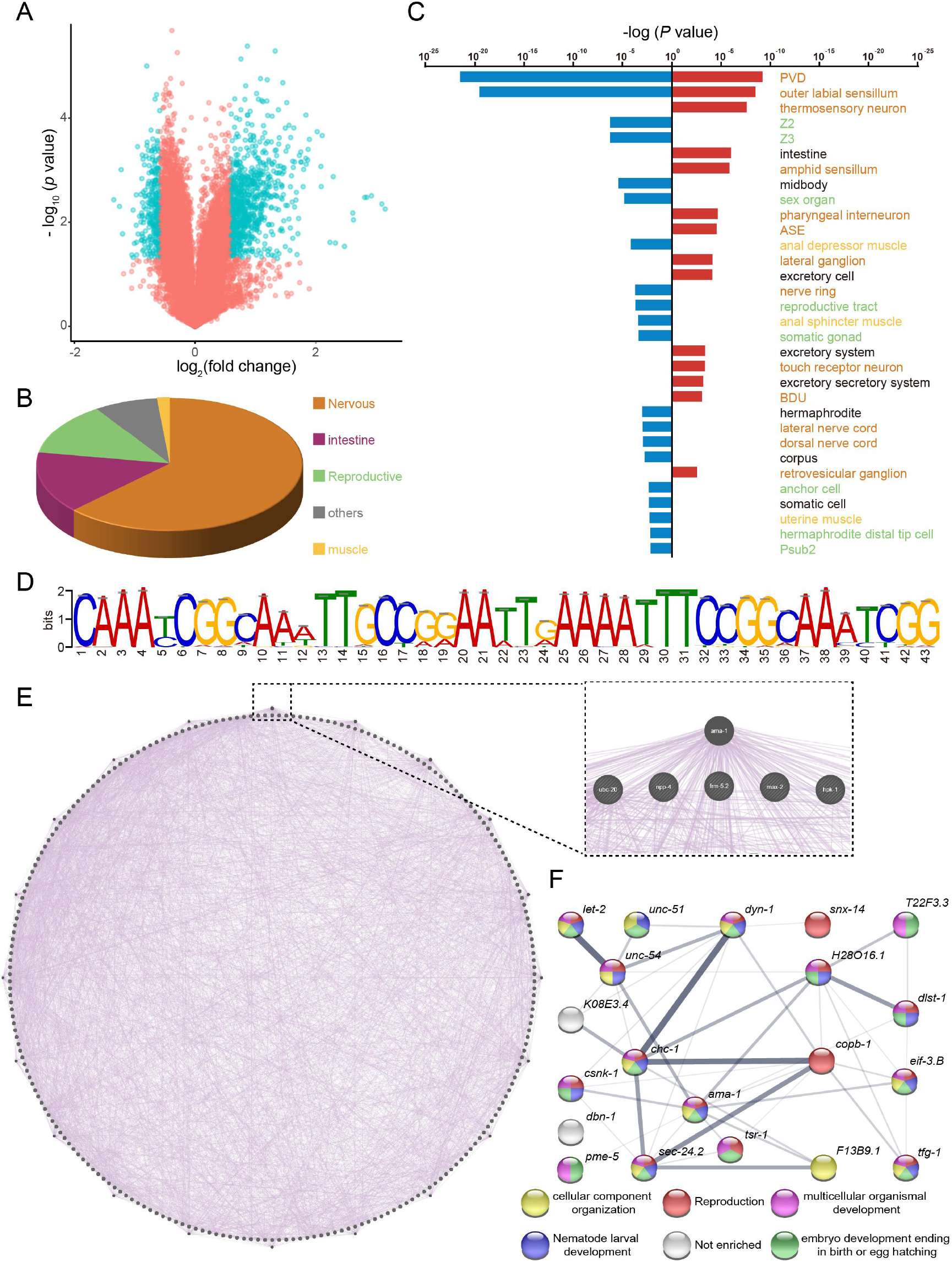
The nervous system is the most affected organ in *C. elegans* after gold NPs exposure. (A) Volcano plot depicting changes in gene expression in *C. elegans* after gold NPs exposure was analyzed. Gene fold changes are plotted against a *P* value. Genes with upregulated or downregulated by more than 1.5 fold and with a FDR less than 0.05 are marked as blue circles. Genes with upregulated or downregulated by less than 1.5 fold and with a FDR more than 0.05 are marked as red circles. (B) Pie graph and (C) bar graph of the tissue enrichment analysis of genes upregulated (blue) and downregulated (red) in gold NPs exposed *C. elegans* group. (D) DNA motif sequence enriched in the promoter region of genes upregulated in *C. elegans* after gold NPs exposure. (E) The protein-protein interaction map of genes upregulated in the gold NPs exposed C. elegans group. (F) The protein-protein interaction between the top 20 upregulated genes. The biological process of each gene was marked based on gene functions.

### Gold NPs regulate development and reproduction through the nervous system

To investigate how gold NPs interact with *C. elegans* in the culture environment, the size distribution and zeta potential were measured in water and S medium. With the increase of concentration, the real size of gold NPs increased (Fig.2A&2B). The average hydrodynamic diameter of 1, 10, 100 mg/L gold NPs in S medium are 94.4 ± 29.9, 952.4 ± 150.1, and 1144.3 ± 159.7 nm, respectively. There was no significant difference between the zeta potential of gold NPs in water and in S medium. The zeta potential for S medium are −5.93, −14.6, and −6.6 mV respectively (Fig.2C&2D). These results indicated that gold NPs have a larger size in S medium than water, but the dispersion is very stable after sonication. To further understand how gold NPs affect the function of the nervous system, gene ontology analysis was proceeded by the list of nervous system enriched genes. From the 32 significantly enriched terms, 10 of them were related to development, including larval, embryo, genitalia, multicellular organism, vulval, gonad, germ cell, and dendrite development. Several cell signaling pathways were shown to be affected, as endocytosis, apoptosis, phosphorylation, cell differentiation, secretion. There are also some phenotype related changes as reproduction, locomotion, morphogenesis, oviposition, lifespan, oogenesis and oocyte maturation (Fig. 2E). These results indicated that Gold NPs exposure regulates development and reproduction through the nervous system. To further confirm the effect of gold NPs on the development of *C. elegans*, the developmental assay was preformed by start exposure from L1 stage. As shown in Fig.2F and 2G, expose to 10, 100 mg/L of gold NPs significantly affected the development of *C. elegans.* The body length of control, 1, 10, 100 mg/L gold NPs are 1068.2 ± 95.3, 1086.6 ± 108.3, 1028.2 ± 155.8, and 771 ± 86.9 μm respectively. After 60 hours of culture, the wildtype control group have 41.4%of worm developed to adult, 55.2% of worm as young adult, 3% as L4 stage. The 1 mg/L gold NPs have 44.1% adult, 50.4% as young adult, 5.5% as L4 stage. For the 10 mg/L exposure group, 44.0% adult, 26.7% as young adult, 29.3% as L4 stage. For the 100 mg/L exposure group, 0.8% young adult, 92.3% as L4 stage, 6.8% as L3 stage (Fig.2H). These results indicate that the 10 and 100 mg/L gold NPs delayed the development of *C. elegans* in a concentration dependent manner.

**Figure 2.**
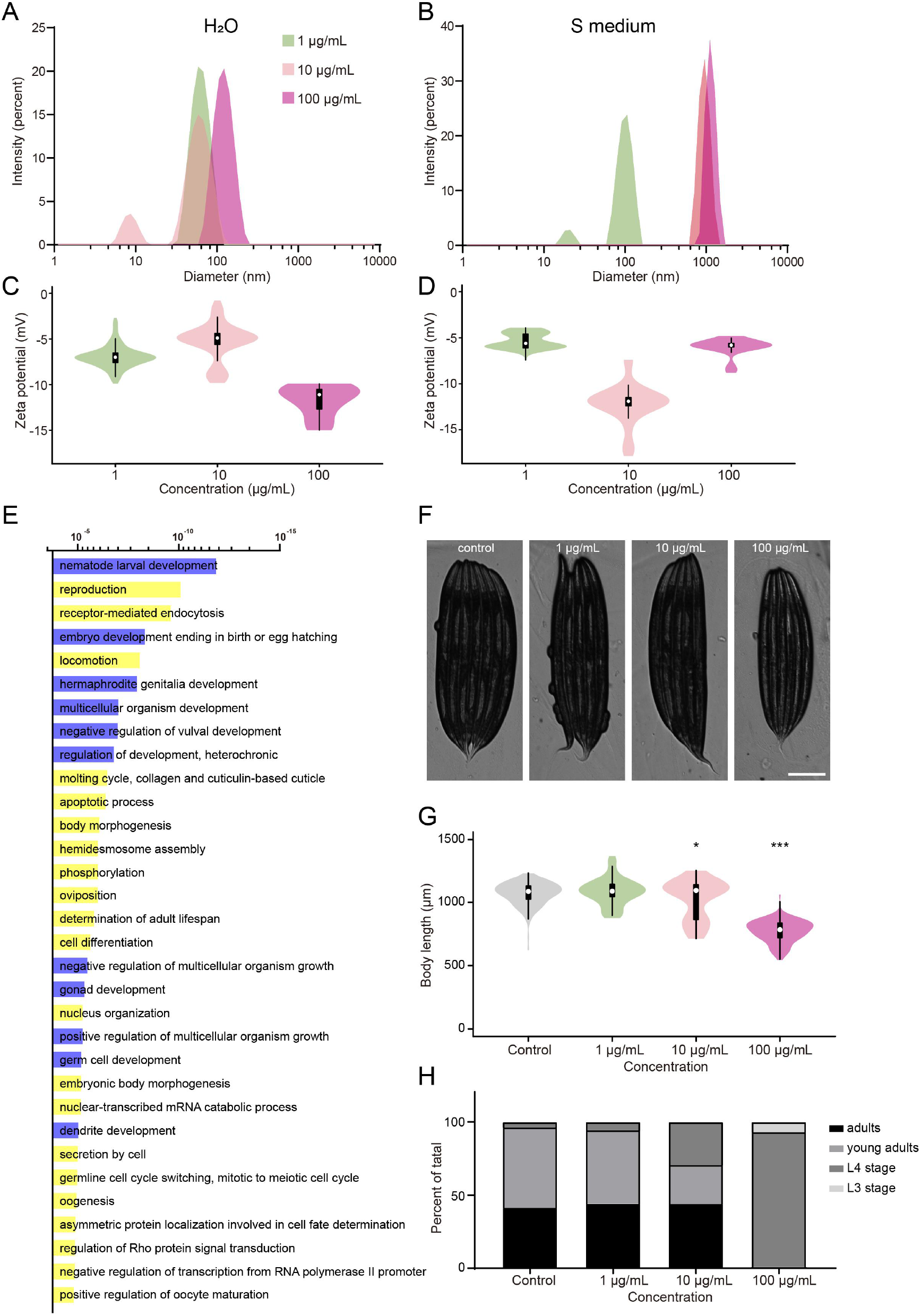
The developmental stage analysis of the gold NPs exposed C. elegans group. The size distribution of 40 nm gold NPs in water (A) and S medium (B). The zeta potential of 40 nm gold NPs in water (C) and S medium (D). (E) Bar graph of the biological process analysis of genes upregulated in the gold NPs exposed *C. elegans* group based on *P* value. The development related biological process was marked as blue, others was marked as yellow. (F) The images of L1 worm exposed to 0, 1, 10, 100 mg/L gold NPs after 60 hours. (G) The quantification of (F). (H) The developmental stage analysis of L1 worm exposed to 0, 1, 10, 100 mg/L gold NPs after 60 hours.

### Gold NPs damage the nervous system mainly through oxidative stress

Since the nervous system was the most affected organ by gold NPs exposure, we further investigated the mechanism of how gold NPs affect the nervous system function. Oxidative stress has been reported as the main factor causes damage when the cells and mice were exposed to gold NPs ^17,19^. We analyzed our data by comparing the reactive oxygen species signaling in *C. elegans* dataset (GSE54024) ^31^. From all the genes significantly increased and enriched in the nervous system, 26.5% of them were related to reactive oxygen species signaling pathway (Fig. 3A). Gene ontology showed these 62 genes were related to development, reproduction endocytosis, locomotion and egg hatching (data not shown). In *C. elegans*, the transcription starts from the transcription factor binding site by recognizing specific DNA motif. DAF-16 is one of the most well-known oxidative stress regulation transcription factors in *C. elegans* ^32^. The motif in the promoter region that DAF-16 binds to have been well established. From the DNA sequence analysis near the promoter region, 59 out of 62 genes contain DAF-16 binding motif. The binding motif in the promoter region of *unc-11, nep-17* and *acs-13* have been graphed (Fig. 3B). To further confirm the activation of insulin-like receptor pathway, *daf-16p::daf-16a/b::GFP* mutant was used. As shown in Fig.3C&D, the worms from 100 mg/L gold NPs exposure group have obverse nucleus localization compare to control worms. The average number of control, 100 mg/L gold NPs, and heat shock group for DAF-16 nucleus localized cells are 10.2 ± 4.5, 48.3 ± 10.7, 164.0 ± 8.6, respectively. To confirm the daf-16 regulated oxidative stress respond genes were activated by gold NPs exposure, we preformed the real time PCR analysis. The enriched genes as *cpr-5, sop-2*, and *acly-1* are not significantly changed in mRNA expression. While genes as *nkcc-1, lpr-3*, and *mex-5* are significantly increased in mRNA expression. The difference in the RNA-seq enriched target genes and our real time PCR results might due to the difference in worm culture medium, and the size and synthesis method of gold NPs. This result indicates that the nervous system regulates gold NPs exposure induced oxidative stress mainly through transcription factor DAF-16. The oxidative stress downstream gene *nkcc-1, lpr-3*, and *mex-5* are the target genes for gold NPs exposure.

**Figure 3.**
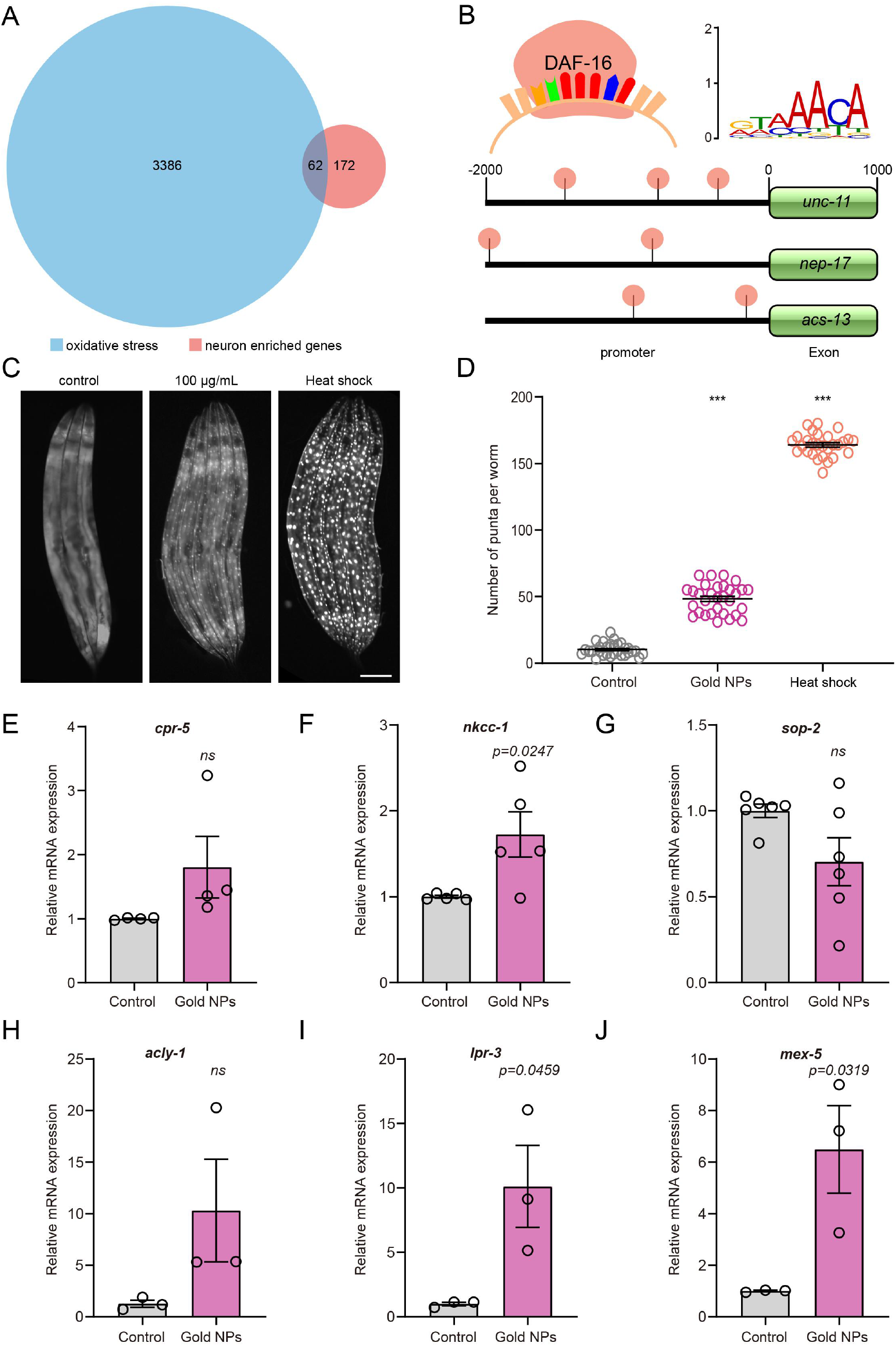
The gold NPs induced oxidative stress in the nervous system were regulated through transcription factor DAF-16. (A) Venn diagram of genes enriched in oxidative stress induced by treatment with the pro-oxidant paraquat and genes enriched in nervous system induced by gold NPs. (B) The location of DNA motif recognized by transcription factor DAF-16 in the promoter region of *unc-11*, *nep-17* and *acs-13*. (C) The image of daf-16 subcellular localization for control, 100 mg/L gold NPs and heat shock groups. (D) The quantification of (C). (E-J) The mRNA expression of *cpr-5, nkcc-1, sop-2, acly-1, lpr-3*, and *mex-5* for control and 100 mg/L gold NPs exposure group.

### Gold NPs enriched oxidative stress genes are mainly expressed in the cytoplasm and nucleus in neuron cells

Since oxidative stress is the main way for gold NPs to damage neuron cells, we further analyzed the subcellular distribution of the significantly increased and nervous system enriched genes that related to the reactive oxygen species signaling pathway. From the protein sequence based subcellular localization prediction, 57 out of 62 genes have been localized to each specific organelle. Forty genes were predicted to be localized in the cytoplasm, 23 in the nucleus and 3 in the mitochondrion (Fig.4A and 4B). Neuron cells communicate with each other by using signal molecules to send and receive ions and transporters. Therefore, changes in ion channel proteins, transporters, receptors, motor proteins and complexes have a huge impact on the stress response, movement and energy metabolism of the organism. Hereby, we analyzed the ion channel genes that significantly changed in expression when worm exposed to gold NPs. The *twk-47* from K+ channel, *ncx-2* and *nkcc-1* from SLC channel, *nep-17* and *nep-18* from neuropeptides metabolism. The *egl-30, gpb-1* and *snx-14* from downstream of GPCRs, the *dlg-1, afd-1* and *shn-1* from PDZ domain proteins, the *klp-7* and *klp-10* from kinesin-like motor proteins, the *hum-7* from myosin motors, the *unc-52* and *mig-6* from extracellular immunoglobulin and leucine-rich repeat domain-containing proteins and the *hmr-1* from cadherins were significantly increased in *C. elegans* exposed to gold NPs. This result indicates that the proteins affected by gold NPs exposure were mostly located in the cytoplasm and nucleus. The communication of neuron cells might be affected through several significantly changed channel proteins and neuron transmitter proteins.

**Figure 4.**
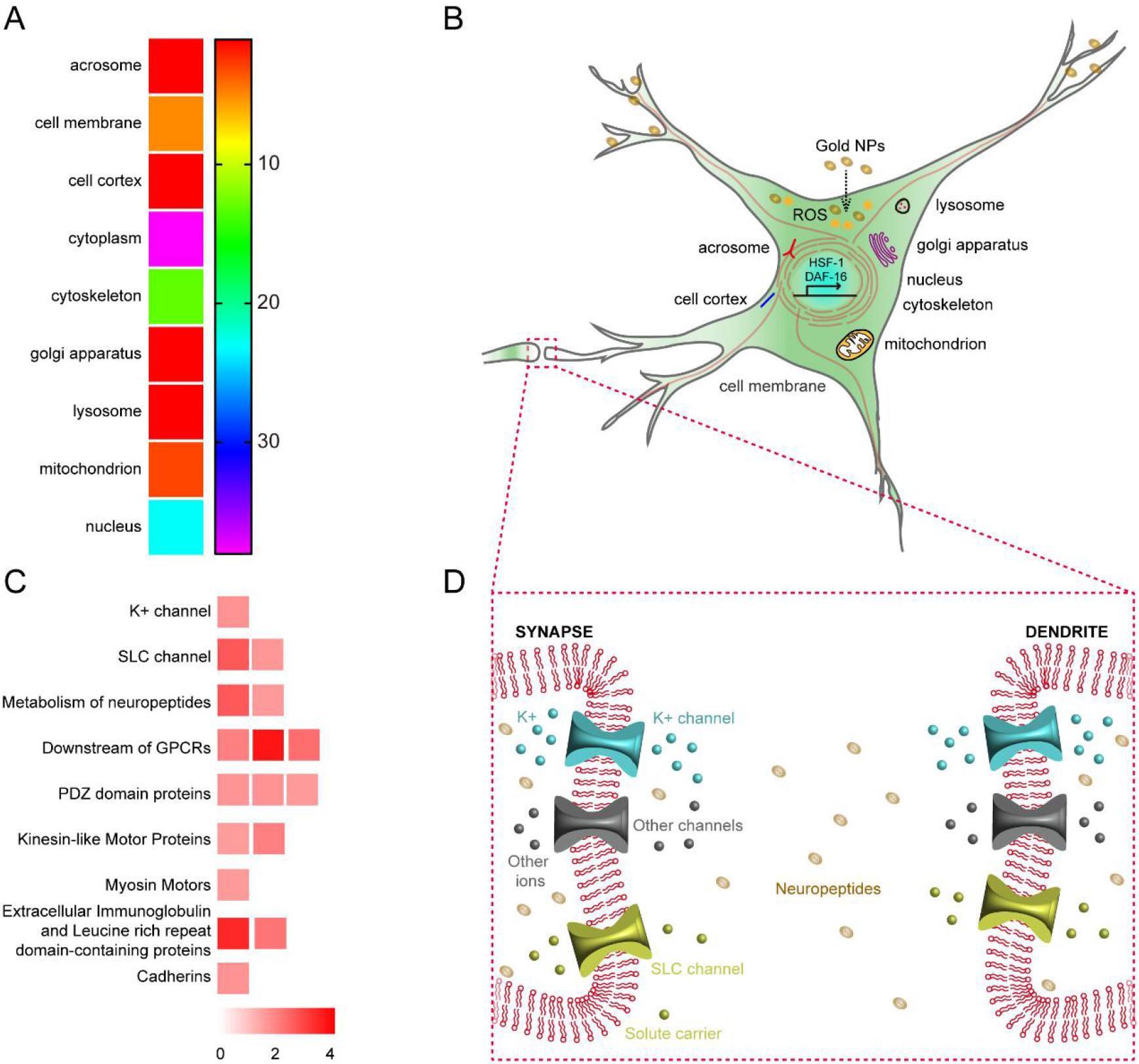
The gold NPs induced oxidative stress affects multiple organelles in neuron cells and their communication. (A) The heat map and (B) the schematic diagram of subcellular location of the oxidative stress genes enriched in neuron cells. (C) The heat map and (B) the schematic diagram of the oxidative stress affected channel genes which functions in neuron cell communication.

## Discussion

Considering the growing interest in gold NPs used in nanomedicine and target drug delivery, the potential health effect of gold NPs need to be well studied. A lot of research has been done to investigate the effect of gold NPs on cancers, the lung and neuron cells ^4,16,33^. More detail needs to be investigated at the organ level. Although biodistribution assays have been done on *C. elegans* and mice ^22,34^, the organ (intestine for worm, liver, kuffer cells and spleen for mice) have most gold NPs accumulated, has a higher possibility to be the organ damaged. However, based on the analysis of whole-body RNA-seq data from *C. elegans*, our research shows the nervous system was most affected.

Many researchers have demonstrated that gold NPs exhibit less cytotoxic and genotoxic effects compared to other nanomaterials like Zn and Ag NPs ^35,36^. However, research in *Drosophila melanogaster* reported the gold NPs induces decreased infertility and numbers of progeny, increased in aberrant phenotype ^37^. These results indicating that gold NPs might affect the development of animals, mostly in the reproductive system. Similar research conducted in *C. elegans* claimed that gold NPs maternal exposure causes a trans-generational effect ^38^. Furthermore, gold NPs disrupt eye development and pigmentation in Zebrafish ^39^. Our finding claimed that most of the nervous system enriched genes, which significantly changed in expression in gold NPs exposed worms, are development related genes. Data from developmental stage analysis also confirmed our finding. Future studies will investigate how neurons regulate development when exposing to gold NPs.

Many cellular responses induced by gold NPs, such as unfolded protein stress, apoptosis and DNA damage, are characterized by an increase in the oxidative stress. Previous experiments demonstrated that 1.4 nm gold NPs induce necrosis in HeLa cells by oxidative stress further amplified by mitochondrial damage ^17^. Gold NPs induced apoptosis and upregulation of pro-inflammatory genes were discovered ^15^. The researcher claimed that smaller Au NPs with larger surface area, might yield higher levels of oxidative stress. Gold NPs with a diameter less than 50 nm can easily penetrate the cell membranes and enter the nucleus of the cells ^33,40–42^. In our data, 40 nm gold NPs significantly increased the mRNA expression of several *daf-16* regulated oxidative stress genes, which showed how the neurons response to the oxidative stress.

Gold NPs modulate neural activity has been studied in multiple nervous systems ^43,44^. It has been shown that gold NPs increase neurite length, promoter adhesion and proliferation, change the action potentials, and affect Ca^2+^ influx ^45–47^. Furthermore, engineered gold NPs were able to bind to sodium channels, transient receptor potential vanilloid member 1 channels, and P2X3 receptor ion channel in dorsal root ganglion neurons, which affect the neuron-neuron communication ^44^. Our data showed that genes related to potassium channel, SLC channel, GPCR, cadherins, motor proteins were upregulated by gold NPs exposure, which indicated that gold NPs might affected the function of related channels. Further research should be conducted on how related channels regulates the ions further affect neuron communication.

## Acknowledgement

We are grateful to our colleagues Cristina Johnson Kay and Nicole Encalada for critical reading of the manuscript.

